# Hierarchical organization endows the kinase domain with regulatory plasticity

**DOI:** 10.1101/197491

**Authors:** Pau Creixell, Jai P. Pandey, Antonio Palmeri, Marc C. Santa-Olalla, Rama Ranganathan, David Pincus, Michael B. Yaffe

## Abstract

The functional diversity of kinases enables specificity in cellular signal transduction. Yet general rules for how the kinase domain allows the more than 500 members of the human kinome to receive specific regulatory inputs and convey information to appropriate substrates – all while using the common signaling currency of phosphorylation – remain enigmatic. Here, using co-evolution analysis and quantitative live-cell assays, we reveal a deep hierarchical organization of the kinase domain that facilitates the orthogonal evolution of regulatory inputs and substrate outputs while maintaining catalytic function. Three quasi-independent functional units in the kinase domain (known as protein sectors) encode for catalysis, substrate specificity and regulation, and these distinct subdomains are differentially exploited by somatic cancer mutations and harnessed by allosteric inhibitors. We propose that this functional architecture endows the kinase domain with inherent regulatory plasticity.

## INTRODUCTION

The ability of cells to specifically respond to a wide variety of environmental cues is made possible by the capacity of signaling proteins to form both insulated and overlapping information-processing networks. Protein kinases are critical nodes in these networks due to their ability to transmit a major signaling currency – phosphorylation – that can modify the activity, localization, interactions, stability and other functions of their substrate proteins^1–4^. As such, kinases have diversified into more than 540 distinct proteins within the human proteome^2^. While by definition all kinases share the core function of phosphorylating substrates, the evident specificity of signaling pathways indicates that kinases have evolved divergent substrate recognition capabilities and regulatory mechanisms. How these evolving kinase domain family members accomplished the balancing act of maintaining catalytic function while accommodating a diverse range of novel substrates and regulatory inputs remains a mystery.

Three features of protein kinases make solving this mystery particularly worthwhile. First, the kinase domain is the domain most often found encoded in cancer genes^5^. Second, there remains a major unmet need to develop allosteric drugs that perturb specific kinases. These drugs will have to take advantage of differences in substrate specificity and regulation rather than acting as ATP mimetics, which often results in off-target effects^4,6,7^. Finally, from a synthetic biology perspective, the kinase domain represents a highly plastic molecular machine that should be able to be programmed to dynamically convert a broad range of molecular inputs to a diverse array of orthogonal outputs^8–10^.

Here, we sought to uncover the functional architecture of the kinase domain. To this end, we developed a computational approach, termed comparative coupling analysis (CCA), to define groups of co-evolving functional residues (protein sectors) in the kinase domain, both at the whole-kinome level and within well-defined kinase subgroups. CCA predicted the existence of three quasi-independent sectors that encode distinct functions: catalysis, substrate specificity and regulation, which we validated via mutational analysis coupled to quantitative live cell measurements. We find that the three sectors display a hierarchy of conservation that corresponds to the functional plasticity that each sector demands. The most conserved sector encodes the fundamental catalytic function required by all kinases, while the sectors that encode substrate recognition and regulatory inputs show progressively less conservation and more subfamily specificity. The sectors are exploited both by cancer mutations and allosteric kinase inhibitors, thus underscoring their functional relevance. Our results indicate a hierarchical organization of the kinase domain by which substrate recognition and regulatory inputs can be readily altered over evolution and tuned by mutations and inhibitors.

## RESULTS

### Three groups of co-evolving residues constitute distinct sectors within the kinase domain

To gain insight into the hard-wired functional organization of the kinase domain, we developed a novel computational approach as an extension of the statistical coupling analysis (SCA)^11,12^.SCA was originally designed to identify groups of amino acids known as protein sectors that coevolve to perform a specific molecular function^11,12^. Our method – termed comparative coupling analysis (CCA) – differs from SCA in that it uses information from protein family subgroups to predict similarities and differences in the sector-forming residues and the conserved and evolving functions the sectors encode (Figure 1A-C).

**Figure 1.**
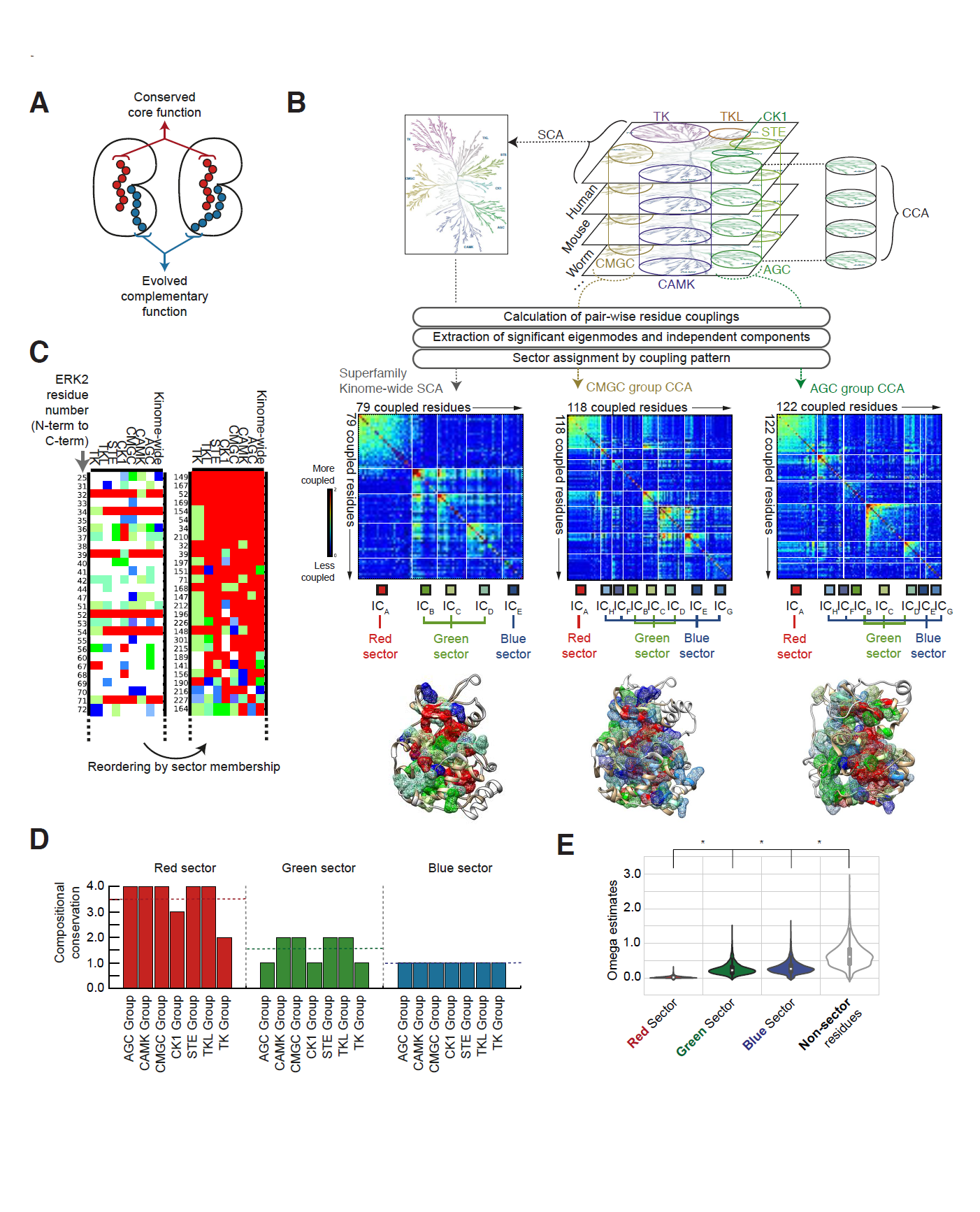
Comparative coupling analysis of the kinase domain. A – Conceptual representation of amino acid residues driving specific molecular functions allows the distinction between those residues involved in “*core”* functions (such as catalysis, shown in red), which will be highly conserved in all instances of that domain, and those residues involved in “*complementary”* functions (like regulation, shown in blue), with higher freedom to evolve. B – Left: In Statistical Coupling Analysis (SCA) of protein kinases, a superfamily-wide domain alignment from a single species is used to calculate coupling between every pair of residues and every pair of positions in the alignment. The resulting high-dimensional matrix is further compressed into a two-dimensional coupling matrix, where the value at every position represents the degree of coupling between every pair of positions in the domain. From this coupling matrix, significantly coupled residues are identified by spectral decomposition and comparison to a randomized alignment in which residues are randomly reshuffled at each position in the alignment, thereby maintaining conservation while losing coupling. Next, independent component analysis (ICA) allows the identification of “independent components” (ICs) or clusters of residues that are more coupled amongst themselves than with the other residues. Finally, in cases where significant coupling still exists between several “independent components”, they are considered part of the same protein sector (see Methods). Right: Unlike SCA, where kinases from all groups are considered at once, for Comparative Coupling Analysis (CCA) subgroup-specific alignments are performed using ortholgs from 15 divergent genomes where each of the seven classically defined protein kinase subgroups^2^ (TK, TKL, CK1, CMGC, STE, CAMK and AGC) is considered individually. This subgroup-specific analysis typically identifies both a larger number and a distinct set of coupled residues as shown for AGC and CMGC. A full representation of all CCA results for the each of the different kinase subgroups is shown in supplemental figure S1. C – Representation of a fraction of all coupled residues within the kinase domain colored by their independent component and sector membership in the different groups ordered by primary sequence (left) or by sector membership after clustering (right). For a full representation of all coupled residues, see supplemental figure S1. D – Quantification of compositional conservation for the different sectors as measured by the median number of other kinase subgroups for which a red sector residue (or green or blue) is also red (or green or blue). E – Estimation of the degree of negative selection for the different protein sectors identified within the kinase domain by calculating omega estimates corresponding to the number of synonymous and nonsynonymous substitutions for the different residues, while correcting for multiple substitutions, transition/transversion rate biases and base/codon frequency biases^13^ (see Methods).

We applied both SCA and CCA to the kinase superfamily, which consists of seven canonical, well-defined subgroups classified on the basis of function, sequence and structural similarity, evolutionary history and substrate class (AGC, CAMK, CMGC, STE, CK1, TKL and TK)^2^. Sequences were obtained for 4867 kinases covering all seven kinase subgroups from fifteen different species ranging from Homo sapiens to Giardia, and aligned using the alignment of human kinase domains as a reference (Table S1, see methods). SCA performed on the entire collection of aligned sequences identified five independent components in the kinase domain that collapse to three distinct sectors, herein referred to as the red, green and blue sectors (Figure 1B). CCA revealed that the three sectors identified in the kinome-wide alignment are all present in every kinase subgroup (Figure 1B). However, importantly, CCA revealed critical differences in the subsets of the residues that form the sectors and in the degree of conservation of the sectors among the seven different kinase subgroups (Figure 1B–C, Figure S1 and Table S2). Indeed, while the red sector is compositionally conserved, meaning that the same residues largely comprise the red sector in all subgroups, the green and blue sectors show progressively less compositional conservation (Figure 1C–D). To investigate potential evolutionary differences between the three protein sectors, we estimated the number of nonsynonymous substitutions that occurred in residues forming the different protein sectors (Figure 1E and methods)^13^. The significantly different number of nonsynonymous substitutions estimated for the three sectors suggests that they are distinct evolving units resulting from diverging evolutionary pressures.

### The red sector includes the conserved catalytic core of the kinase domain

The red sector encompasses known kinase architectural determinants that are important for catalytic transfer of phosphate from ATP to a substrate hydroxyl group (Figure 2A and Figure S2). These determinants include glycine residues in the P-loop, the DFG and APE motifs that delimit the activation segment, the catalytic loop HRD motif and parts of the catalytic and regulatory spines^1–3,14,15^ in addition to other residues that co-evolve with them and whose direct contribution to catalytic function has been less well recognized. Indeed, consistent with the core function of this sector, we observed a direct correlation between primary sequence conservation of a given residue and its contribution to the red sector (Pearson’s correlation of 0.79, Figure 2B); a direct correlation that was not present for other sectors (Figure S2). The observation that specific core residues involved in catalysis were contained in the red sector, however, was perplexing, since strictly conserved residues that do not show co-variation with other amino acids should be invisible upon SCA and CCA analysis. Therefore, to clarify this and identify which specific kinases drove the identification of this sector, singular value decomposition (SVD) was performed. SVD analysis revealed a group of known catalytically-impaired pseudo-kinases^16^ as those that presented sequence variations in the red sector (Figure 2C). Taken together, these results suggest that the red sector represents coupled residues that delineate the deeply conserved catalytic core of the kinase domain.

**Figure 2.**
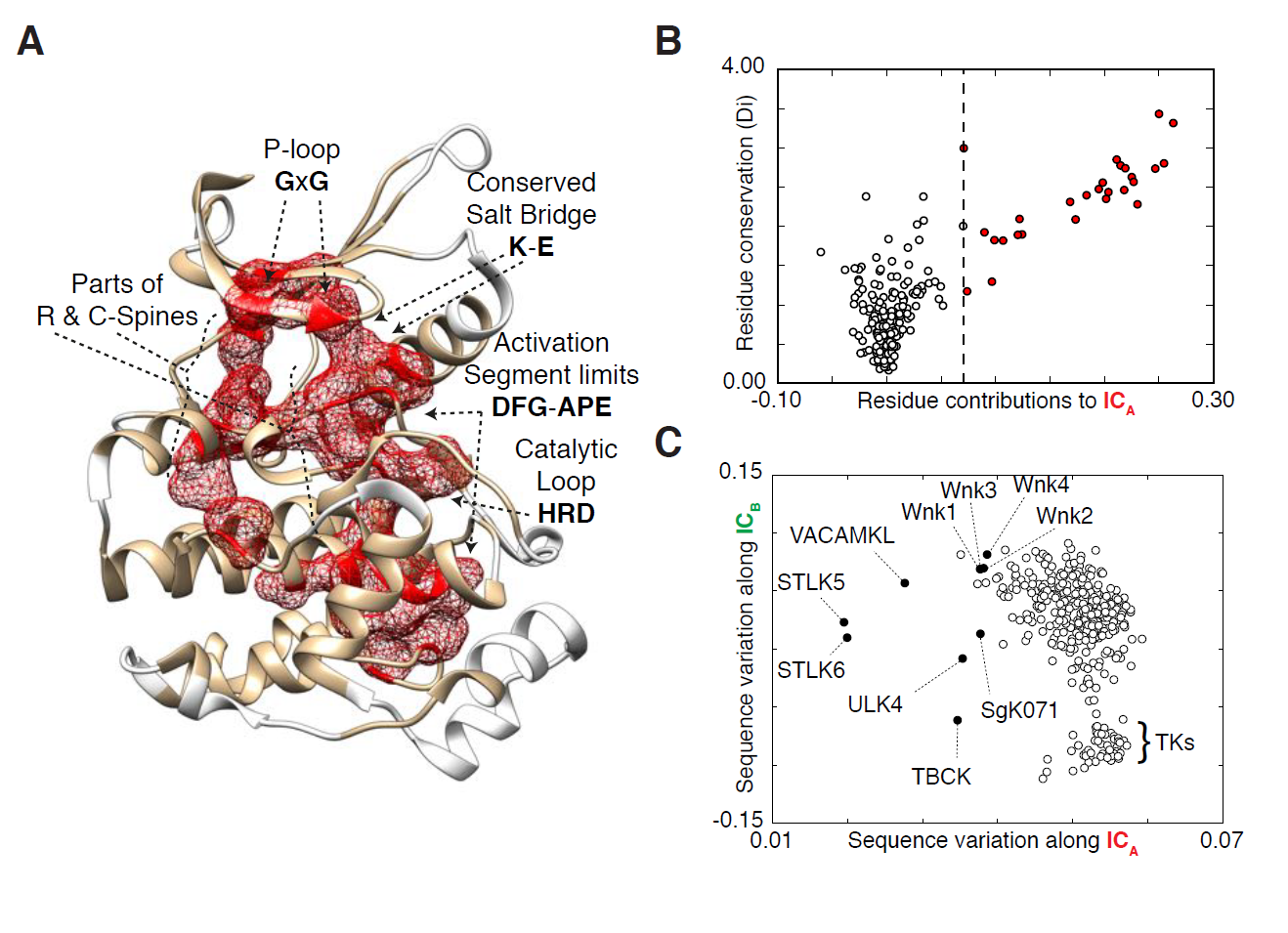
The red sectors drives catalysis. A – Depiction of residues contained within the red sector, illustrated on the structure of ERK2 (PDB: 4QTE). Residues that have been previously established as critical components of the catalytic core of the kinase domain are indicated. B – Comparison between the contribution of every residue within the kinase domain to the first independent component (the one constituting the red sector) and the degree of overall conservation of that same residue. Residue conservation is measured using the Kullback-Leibler relative entropy, Di (see Methods). C – Scatterplot positioning protein kinases according to their sequence variation along the first and second independent components, as described in Figure 1B. The group of pseudo-kinases, which form the majority of kinases diverging along the first independent component (the red sector), are shown in filled black circles. Notably, the second independent component, part of the green sector, separates the tyrosine kinases (TKs).

### Green sector composition determines substrate specificity

The green sector displays intermediate compositional conservation (Figure1D). The green sector is formed by residues that line and bracket the substrate binding site, and includes known determinants of substrate specificity such as the P+1 loop and residues downstream of the HRD motif near the catalytic loop (Figure 3A) ^1,3,17^. Indeed, in several kinase-substrate co-crystal structures, the green sector is the sector that makes the largest amount of direct substrate contact, as illustrated by the structure of AKT/PKB in complex with GSK3 (Figure 3B) and the structure of PKA in complex with PKI (Figure S3). Further supporting a role in substrate recognition and specificity, SVD revealed that the three independent components that make up the green sector broadly separate kinases on the basis of their substrate specificity: tyrosine kinases (TKs) are clearly separated from the rest of the kinome, which is comprised of serine/threonine and dual-specificity kinases (Figure 2C, 3C). Moreover, SVD organized the non-TK kinome along a substrate specificity gradient, ranging from the proline-directed CMGC kinases toward basophilic-directed CAMK and AGC kinases (Figure 3C, upper right and lower left, respectively). As an orthogonal approach, we compared the KINspect score^17^ – an established metric to quantify the likelihood that a residue has a role in substrate specificity – for each green sector residue across the kinase domain to the score for all non-green sector residues. Green sector residues showed a significantly higher KINspect score than non-green sector residues (Mann Whitney U, p = 0.005, Figure 3D). Taken together, these results suggest that the green sector is composed of coupled residues with functional roles in substrate recognition and specificity.

**Figure 3.**
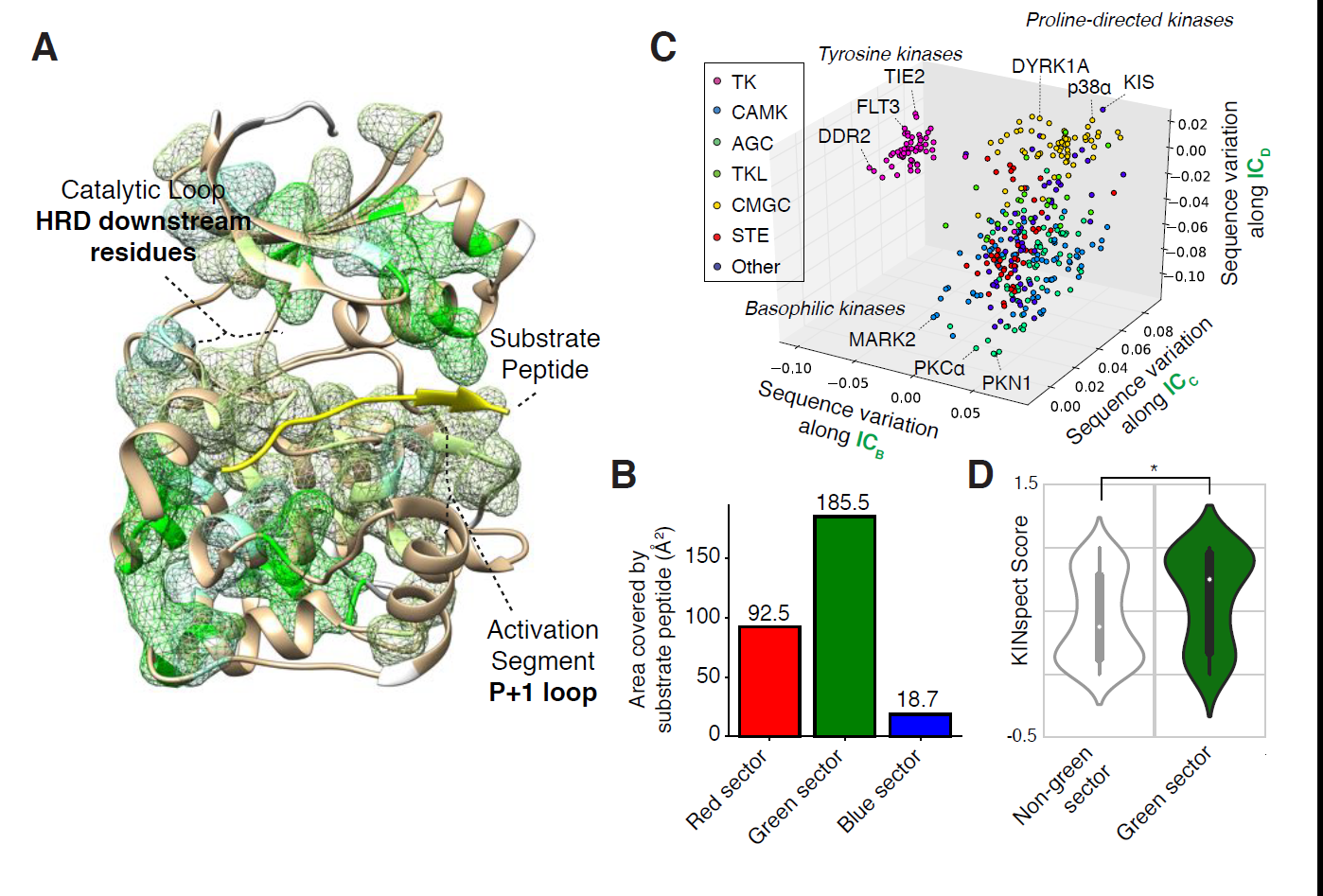
The green sectors encodes substrate peptide specificity. A – Illustration of residues contained within the green sector, superimposed on the structure of PKB/AKT in complex with a GSK3 peptide substrate with sequence GRPRTTSFAE (PDB: 4EKK). Regions previously implicated as key determinants of substrate specificity are indicated. B – Barplot comparing the surface area of red, green and blue sector residues that is buried by the peptide substrate in the structure of the PKB/AKT:GSK3 substrate peptide complex, shown in panel A. For this calculation, the solvent exposure of all residues is calculated in the presence and absence of the peptide substrate and the differencial exposure for the different sector residues is displayed. C – Scatterplot of sequence variation between all human kinases, relative to one another, of residues that form the second, third and fourth independent components as described in Figure 1B. Each point indicates a particular kinase and is colored according to its major kinase group. D – Violin plot of the distribution of KINspect scores, an orthogonal measure of the contribution of each residue within the kinase domain towards substrate specificity^17^, for residues belonging to the green sector compared to residues outside the green sector. The width of the violin at any particular KINspect score indicates the number of residues that match that score.

### The blue sector is poised to receive regulatory inputs

The most highly divergent sector between subgroups, the blue sector, appears to connect the active site and residues in the red and green sectors with more peripheral sites at the surface of the kinase domain (Figure 4A). This topology naturally suggests a role in transmitting regulatory inputs. Indeed, we found clear examples where regulatory interactions occur preferentially with blue sector sites, such as an allosteric binding site on the yeast MAPK Fus3 for the scaffold protein Ste5 (Figure 4A-C) or the binding site on CDK2 for cyclin A (Figure S4). Taken together, this architectural and anecdotal evidence suggests that the blue sector is well poised to receive regulatory inputs; a hypothesis that we subsequently tested experimentally.

**Figure 4.**
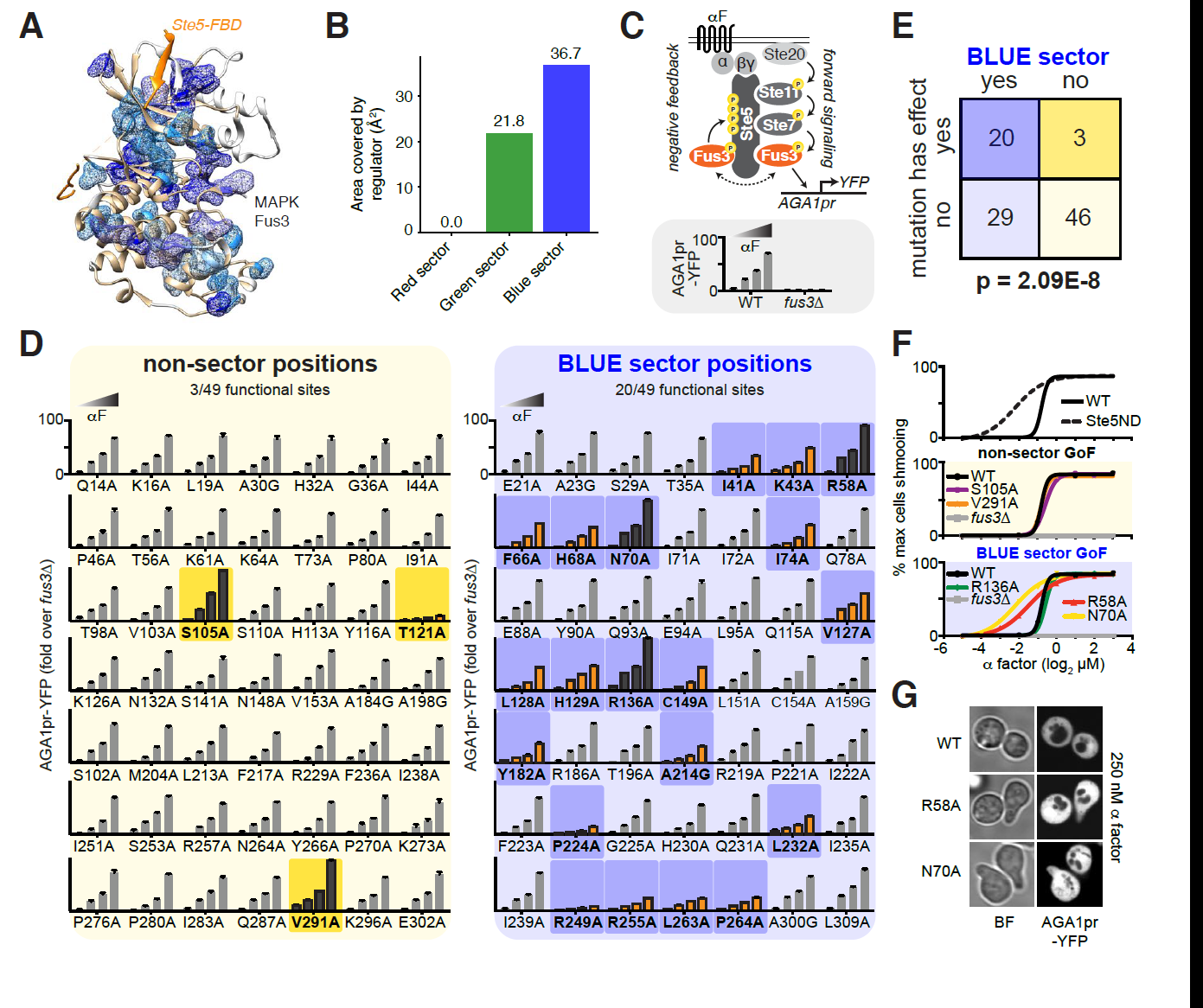
Mutational analysis of Fus3 reveals a functionality for the blue sector in mediating regulatory inputs. A – Illustration of residues forming the blue sector, superimposed on the structure of the yeast MAP kinase Fus3 in complex with the Fus3-Binding Domain (FDB) domain of Ste5 colored in orange (PDB: 2JED). B – Barplot displaying the differential solvent exposed area surfaces for red, green and blue sector sites in Fus3 in the presence or absence of the Fus3-Binding Domain (FBD) of Ste5, as described in the caption of figure 3B. C – Schema of the yeast mating pathway including the role of Fus3 and its allosteric regulation by Ste5-FBD to maintain the pathway in an inactive state. The downstream transcriptional reporter used in the functional assays for Fus3 activity shown in panel D is indicated (top). At the bottom, control experiments demonstrating the Fus3- and dose-dependency of the fluorescent signal by the reporter. D – On the left, fluorescence signal for the four doses of α-factor upon alanine-scanning of non-sector mutants, highlighting gain-of-function (GOF) mutants and loss-of-function (LOF) mutants in black and orange, respectively. On the right, fluorescence signal for the four doses of α-factor upon alanine-scanning of 49 blue sector mutants, again highlighting GOF and LOF mutants in black and orange, respectively. Phenotypic mutants are highlighted by a darker background for the non-sector screen (yellow background) and the blue-sector screen (blue background). E – Statistical significance for the enrichment in phenotypic mutants when mutating sector sites as compared to non-sector sites. F – At the top, regulatory defects in the allosteric regulator Ste5, as illustrated for the case of a non-docking mutant, lead to more graded mating dose-responses as quantified by the percentage of “shmooing” cells. In the middle, non-sector GOF mutants display mating dose-responses that are comparable to wild-type. At the bottom, in contrast to GOF non-sector mutants, two out of the three GOF blue sector mutants display more graded mating dose-responses phenocopying the effects observed for non-docking Ste5. G – Depiction of “shmooing” cells for the Fus3^R58A^ and Fus3^N70A^ in conditions that do not elicit “shmooing” in wild-type cells.

### Mutational analysis validates the functionality of the blue sector

To investigate the functionality of the blue sector experimentally, we performed a comprehensive mutational analysis on Fus3 and employed quantitative activity assays in live cells. Fus3 is specifically activated in response to mating pheromone (α factor) and coordinates cell cycle arrest with the transcriptional and morphological responses required for mating^18–20^. We alanine scanned all 49 residues comprising the blue sector along with 49 non-sector positions evenly distributed along the Fus3 primary sequence. We genetically integrated each of these mutants as the only copy of Fus3 in the genome and assayed for Fus3 activity in response to different concentrations of α factor using a fluorescent reporter of the mating pathway (Figure 4C). In our strain background, reporter output depends strictly on Fus3, and wild type Fus3 (WT) produces a graded α factor dose response (Figure 4C). In this assay, 3/49 non-sector mutants were distinguishable from WT (1 loss-of-function (LoF) and 2 gain-of-function (GoF) phenotypes) (Figure 4D), as defined by being statistically different in at least two doses of mating pheromone. By contrast, 20/49 of the blue sector mutants had altered activity compared to WT (17 LoF, 3 GoF), revealing a significant enrichment of functional sites in the blue sector (Fisher’s test, p = 2.09E-8, Figure 4D, E).

### Blue sector mutants phenocopy disrupted allosteric regulation

The functionality of the blue sector suggests that it may be a conduit for regulatory inputs. If natural regulatory interactions evolved to exploit the blue sector, then specific blue sector mutations should phenocopy the disruption of cognate regulation. To test this, we repurposed our GoF Fus3 mutants and performed additional functional assays. In addition to being regulated by its upstream MAPKK through canonical activation loop phosphorylation, Fus3 is regulated allosterically – both positively and negatively – by the scaffold protein Ste5^18–21^. These dual modes of allosteric regulation allow the cell to simultaneously achieve a graded transcriptional response and a switch-like morphological response as a function of α factor concentration (Figure 4C). The feedback domain (FBD) on Ste5 mediates the negative regulation of Fus3 required for the switch-like morphological transition that leads to formation of the mating projection, or “shmoo”^21^. As such, disruption of the FBD results in graded shmooing across a dose response of α factor (Figure 4F)^21^. We hypothesized that some of the GoF Fus3 blue sector mutants may have lost the ability to be negatively regulated by Ste5. Consistent with this idea, while both of the non-sector GoF mutants retained switch-like shmoo responses, 2/3 of the blue sector GoF mutants showed graded shmooing (Figure 4F, G). Thus, mutation of blue sector residues recapitulated a phenotype associated with impaired allosteric regulation.

### CCA reveals private (subgroup-specific) sector residues

CCA not only validated the presence of the red, green and blue sectors in each of the kinase subgroups, it also indicated that there are potentially important differences in the composition of the protein sector among the subgroups. In other words, whereas some sector sites are shared by several or all kinase subgroups, others are subgroup-specific (Figure 1C). We hypothesized that these subgroup-specific sector residues, or “private” sites, would be enriched for functionality in members of the subgroup in question but not in out-groups. To test this, we first identified AGC and CMGC kinases as the two subgroups predicted to be the most functionally divergent (Figure 5A and S5). SVD revealed that the dissimilarity between AGC and CMGC sector composition was largely driven by eight AGC-specific private sector sites that are non-sector sites in CMGC kinases (Figure 5B).

**Figure 5.**
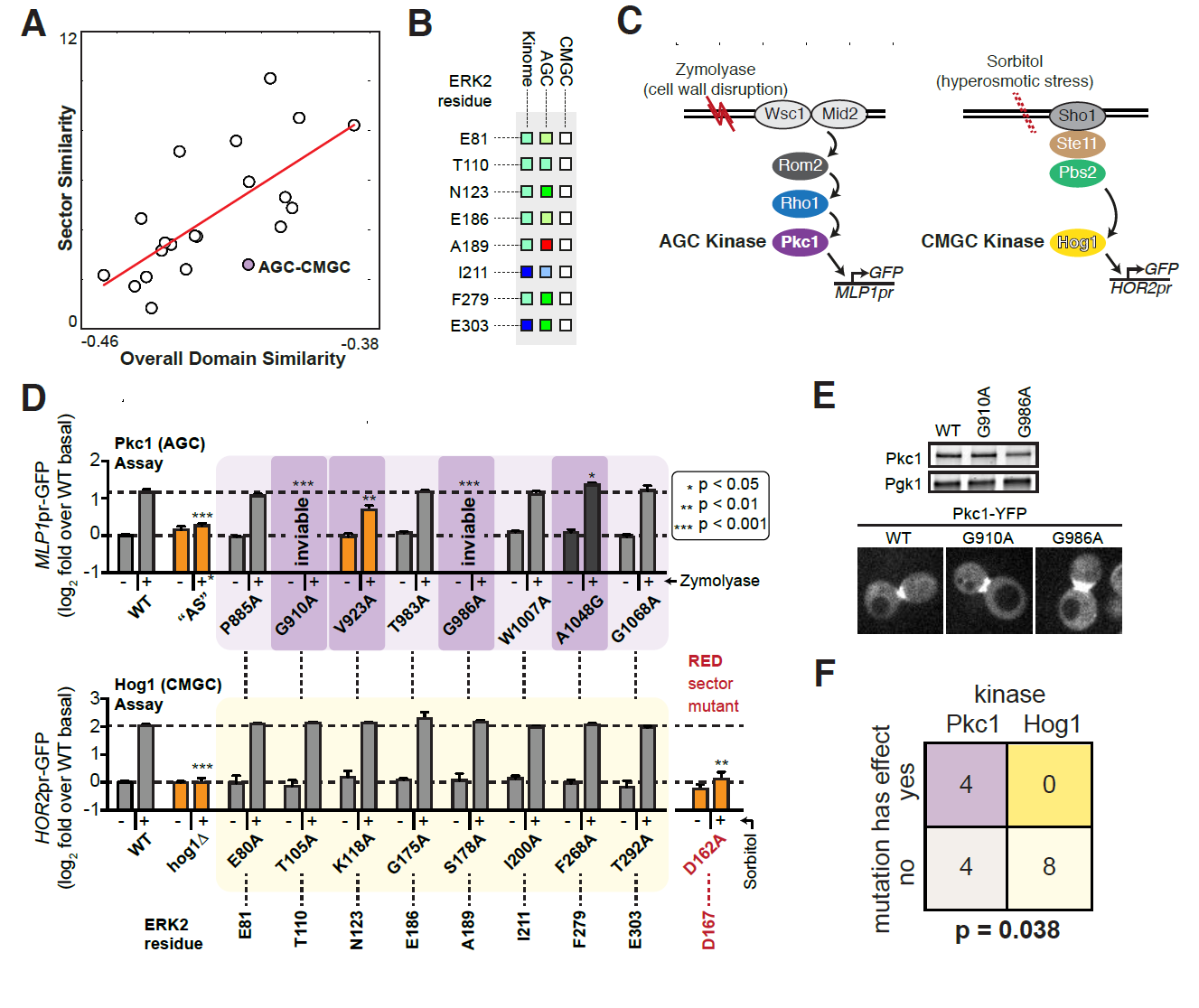
Private sector sites encode subfamily-specific functions. A – Scatterplot including all pairs of kinase groups comparing their overall kinase domain similarity, as measured by overall BLOSUM distance^28^, in the X axis to degree of overlap in their sectors in the Y axis (see Methods). Between the AGC and CMGC subgroups, the residues that define the sectors show limited overlap, despite a moderately high degree of overall similarity between the kinase domain sequences of AGC and CMGC family members. B – Sector memberships of the eight sites that are are part of a sector in the kinome-wide SCA and in the AGC-specific CCA but not in CMGC-specific CCA. These sites are predicted to drive the functional divergence between AGC and CMGC kinases. The amino acid numbering shown corresponds to these sites in the representative structure used to map all CCA models, namely ERK2 (PDB: 4QTE) as described in the methods section. C – Left: Functional assay for Pkc1 where the MLP1-driven downstream transcriptional reporter is activated upon the addition of zymolyase and the subsequent cell wall lysis. Right: Functional assay for Hog1 where the HOR2-driven downstream transcriptional reporter is triggered by hyperosmotic stress resulting from the addition of sorbitol. D – Reporter signal for Pkc1 (top panel) and Hog1 mutants upon mutating the eight AGC-specific private sector sites in both kinases. Analog-sensitive Pkc1 (“AS”) in combination with the analog-specific inhibitor 1NM-PP1 (marked with +*) and Hog1 deletion (hog1D) were used as positive controls for the two assays respectively. A mutant in the aspartate of the DFG motif forming the red sector of Hog1 was used as point mutant control to confirm the Hog1 assay sensitivity to loss-of-function mutants of Hog1. Similarly as in Figure 4D, phenotypic mutants are highlighted by a darker background for Pkc1 screen (darker brown background). No phenotypic mutant was found for any of the eight Hog1 mutants (yellow background). E – Protein expression and cellular localization of wild-type Pkc1, Pkc1^G910A^ and Pkc1^G986A^ as assayed by western blot and microscopy. F – Fisher’s test results assessing the enrichment of phenotypic mutations in Pkc1, as compared to Hog1, upon mutation AGC-specific sector positions (data from Fig.5E).

To experimentally test these AGC-specific sites, we performed mutational analysis on Pkc1, an essential AGC kinase in yeast (homolog of protein kinase C) required for cell wall integrity, and on Hog1, a CMGC kinase (yeast homolog of p38) required for adaptation to hyper-osmotic stress (Figure 5C). Mutation of 4/8 private AGC sector sites on Pkc1 resulted in phenotypic differences compared to wild type: two of the sites were essential for viability while two others altered levels of a fluorescent reporter upon exposure to the cell wall stressor zymolyase (Figure 5D). In contrast to Pkc1, Hog1 tolerated mutations at all 8 AGC-specific sector sites without altered induction of a fluorescent reporter upon addition of sorbitol (Figure 5D). By contrast, Hog1 cannot tolerate a red sector mutation (Figure 5D). Importantly, despite failing to complement for loss of wild type Pkc1, the two inviable mutant proteins were expressed to wild type levels and properly localized when co-expressed with wild type Pkc1 (Figure 5E). These results demonstrate that the private AGC sector residues are specifically and significantly enriched for functionality in an AGC kinase (Fisher’s test, p = 0.038), validating the CCA predictions (Figure 5F).

### Cancer mutations and allosteric inhibitor sites preferentially map onto distinct kinase sectors

Finally, we assessed the relevance of kinase sectors to human disease (Figure 6). Since kinases are key targets of cancer mutations and therapeutic inhibitors^4^, we cross-referenced the sector positions against known somatic cancer mutations (Table S3). After mapping 1,515,599 cancer somatic mutations onto canonical proteins, of which 14,860 mapped onto 13,152 sites within kinase domains, we observed a significant enrichment for cancer mutations at red sector sites (Wilcoxon test, p = 2.4E-4, Figure 6B and methods section). Given that red sector residues comprise the catalytic core of the kinase domain and would thus be predicted to impair catalysis, we were surprised to discover red sector mutations not only in tumor suppressors, but also in kinase oncogenes such as B-Raf and EGFR (Figure 6C, D). Importantly, in addition to mutations involving residues with well-established functions in catalysis (i.e. the DFG motif and Gly-rich loop), this mapping of cancer-associated mutations onto the red sector positions indicates potentially important roles for additional residues that have not been previously implicated in catalytic function. Moreover, in addition to these red sector somatic mutations, we found that allosteric inhibitors of Abl bind at blue sector surface positions, underscoring the regulatory function of the blue sector (Figure 6E). Thus, distinct kinase sectors seem to be exploited by mutations that promote cancer and by small molecule allosteric inhibitors used to control it.

**Figure 6.**
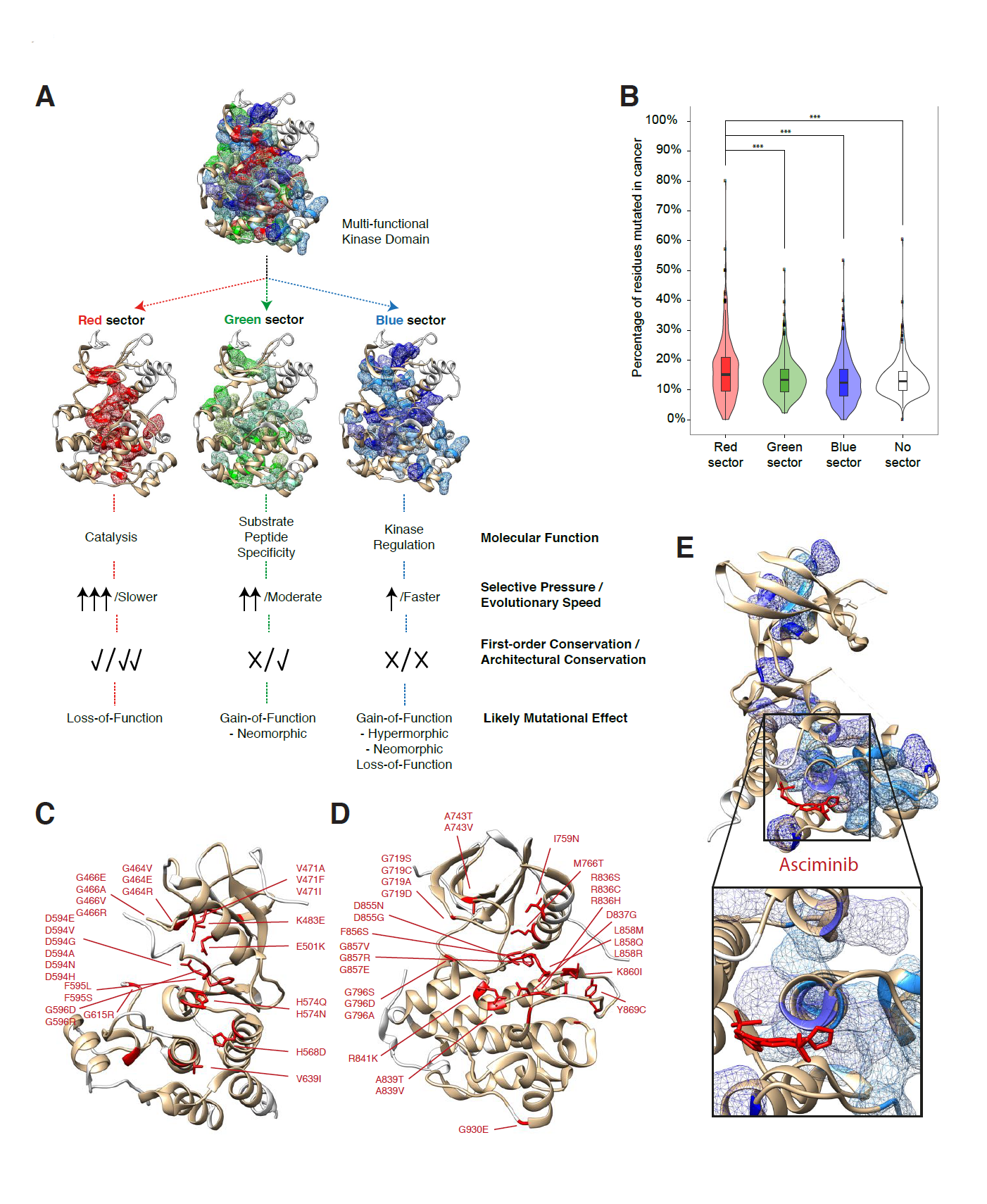
The hierarchical organization of the kinase domain is targeted by somatic cancer mutations and allosteric inhibitors. A – General model of the kinase domain as a highly heterogeneous multi-functional domain with sets of residues encoding distinct molecular functions and being constrained by different selective pressures and evolutionary speeds leading to differential conservation and effects upon mutation. B – Violin plots displaying the distribution of the percentage of residues belonging to the red, green, blue and non-sector sites that are mutated in cancer, using data from the COSMIC repository^29^. Each data point represents a specific protein kinase, for which the percentage of residues in each sector that contain one or more cancer mutations was calculated. p = 5.0E-4; 6.3E-7; 2.5E-4. C – Twenty-seven unique cancer somatic mutations perturbing red sector residues in B-Raf (PDB: 4MBJ). D – Thirty unique cancer somatic mutations perturbing red sector residues in EGFR (PDB: 2GS2). E – Tyrosine kinase (TK) blue sector residues contact the allosteric inhibitor Asciminib in the co-crystal structure of the Abl:inhibitor complex (PDB: 5MO4).

## DISCUSSION

The discovery of distinctly folded domains in proteins led to the interpretation that domains represent units of evolution and function^22,23^. However, it has become clear that there is a large degree of functional and evolutionary heterogeneity among the residues that form a domain. Here we found that the kinase domain is organized in a functional hierarchy that allows a deeply conserved catalytic function to be differentially deployed over evolution to act on distinct sets of substrates and respond to specific regulatory inputs. By formally defining sets of residues likely to encode each of these molecular functionalities using CCA, it becomes tractable to study how these functions may have evolved, or are perturbed in diseases such as cancer (Figures 6).

The residues contributing to the catalytic core of the kinase domain – the red sector sites – are limited in their ability to evolve due to their fundamental importance. Consequently, the red sector is characterized by a high degree of residue conservation and a resulting low degree of evolution. By contrast, the green sector is formed by residues involved in the recognition of the substrate peptide, a molecular function that must – and indeed does – allow plasticity between kinase subgroups to accommodate varied substrates. Finally, the blue sector presents the widest degree of plasticity and divergence between kinase groups, consistent with different kinases evolving divergent regulatory mechanisms. Accordingly, disease mutations can inactivate a kinase via the red sector, but perhaps more sinisterly, could alter substrate specificity or regulatory inputs to subvert and reroute signaling.

There are two notable exceptions to the rule that the red sector is invariant. First, tyrosine kinases represent a functionally divergent subgroup, at least in part driven by their capacity to phosphorylate tyrosine residues instead of serine and/or threonine residues. While the red sector appears to be well defined and compact in all other kinase subgroups, tyrosine kinases present large divergence in all sectors including the red sector. Second, pseudo-kinases also show divergence in the red sector where they have accumulated mutations leading to their impaired catalytic ability. Thus, these exceptions can be easily interpreted and ultimately serve to further support the notion that the red sector drives catalysis.

Beyond the red sector, CCA highlights the functional and regulatory plasticity present in different protein kinases by revealing the variability in the blue sector among the kinase subgroups. At the same time, while CCA allows for this more granular understanding of the architecture of the kinase domain by incorporating subgroup-level information, the *in-silico* identification of functions that are idiosyncratic to individual kinases remains a significant challenge. This limitation arises from the need to capture enough sequences of sufficient diversity. Ultimately, the specific regulatory sites and interactions that control an individual kinase will need to be resolved using focused structural, biochemical and molecular genetic approaches. Despite this caveat, our *in vivo* mutagenesis and functional screens suggest that quantitative experiments performed on a specific protein kinase can recapitulate the functional principles predicted *in silico* at the subgroup-level. After validating the function of these sectors orthogonally, our models provide a means to identify trends and hypothesize mechanisms of action for disease-associated mutations or kinase inhibitors, which can be further tested in focused experiments subsequently.

Our current models still limit the possibility of individual residues performing multiple, overlapping molecular functions. To facilitate interpretation and subsequent analysis, we forced residues to have membership in only a single sector (e.g., residues defined as part of the red sector cannot be simultaneously considered as residues in the green sector). In nature, it is conceivable that a critical residue may play overlapping roles in catalysis, substrate specificity and/or kinase regulation. Similarly, while we define the three sectors as separate entities, there are clear differences in how related to one another the different sectors are. In particular, the blue sector – which connects putative allosteric sites on the kinase surface to the red and green sectors and the active site – includes resides that interface with the other sectors and may contribute to all three functionalities.

The formal, data-driven definition of functional residues presented here enables us to predict the functionality of cancer-associated mutations. We observed enrichment in the number of mutations that perturb red sector sites – not only in tumor-suppressor kinases but also in oncogenes (Figure 6C, D). While initially counterintuitive, this finding suggests that there may be more examples of inactivating mutations leading to roles in trans-activation as has been reported for certain B-Raf mutations^24^–27. Finally, it is exciting to consider that the blue sector residues that we have implicated in kinase regulation appear to serve as portals for allosteric inhibitors (Figure 6E). Targeting blue sector surface sites may lead to the development of next-generation allosteric modulators.

## ACKNOWLEDGEMENTS

We would like to thank Brian A. Joughin, Daniel Lim and all other members of the Yaffe for discussion and critical input leading to this work. We are grateful to Z. Feder and J. Krakowiak for technical assistance, and to the Whitehead Institute FACS facility and the Keck Microscopy facility. This work was supported by a Merck Postdoctoral Fellowship from the Helen Hay Whitney Foundation (to P.C.), an NIH Early Independence Award (DP5 OD017941-01 to D.P.), NIH grants GM104047 and ES015339 (to M.B.Y.), and the Charles and Marjorie Holloway Foundation (M.B.Y.).

## AUTHOR CONTRIBUTIONS

Conceptualization, P.C., D.P., R.R., and M.B.Y.; Methodology, P.C., D.P., and M.B.Y.; Investigation, P.C., J.P.P, D.P., A.P., M.C.S.; Writing P.C., D.P. and M.B.Y. with input from all authors; Supervision, P.C., D.P. and M.B.Y.

## SUPPLEMENTAL MATERIALS

### 5 Supplementary figures

**Figure S1.**
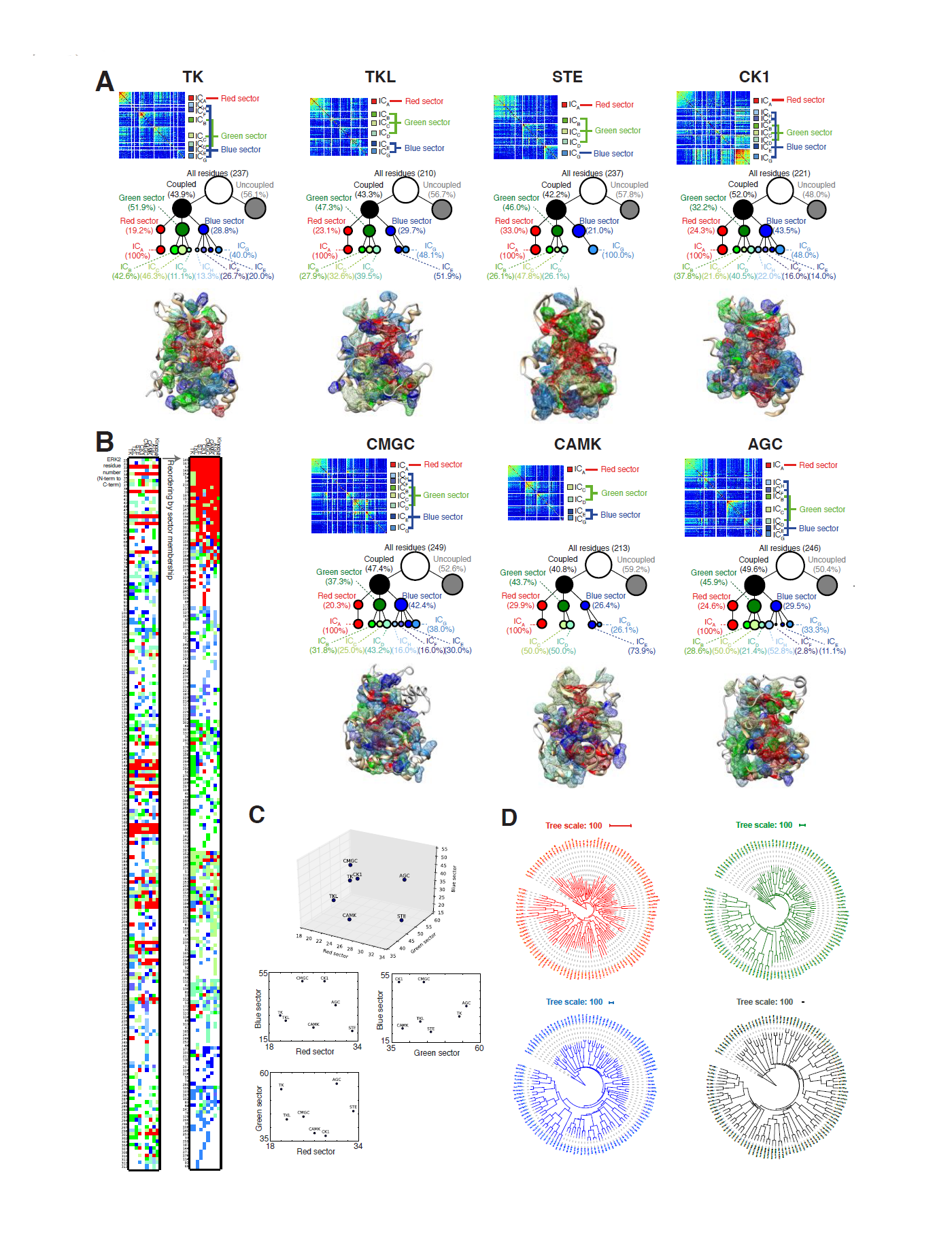
Comparative Coupling Analysis (CCA) computational results. A- For each kinase group, at the top we show the coupling matrix resulting from calculating pair-wise residue coupling, extraction of significant eigenmodes and independent components and identification of protein sectors (as described in the methods section) from each group-specific alignment. In the middle, we represent the percentage of residues that are coupled and within those, the percentage that constitute each sector and independent component. At the bottom, a structural representation displaying the sectors within each kinase group is shown using canonical representative structures.
B- The CCA results from all kinase groups are compared by mapping them back to the largest and most complete structure from all the representatives (ERK2, PDB: 4QTE)^30^. Positions are colored according to their independent component and sector membership in each kinase group. To facilitate interpretation and visualization of the results, we reorder residues based on their sector membership, prioritizing first those residues for which residue membership to red, green or blue sectors is most conserved amongst different kinase groups.
C- Description of the size of the red, green and blue sector for the different kinase groups as a three-dimensional scatterplot as well as with the corresponding three two-dimensional scatterplots.
D- Phylogenetic dendrograms considering only residues that form the red, green or blue sector residues as well as those residues not belonging to any sector. In this case, the CMGC alignment was used to generate the different dendrograms.

**Figure S2.**
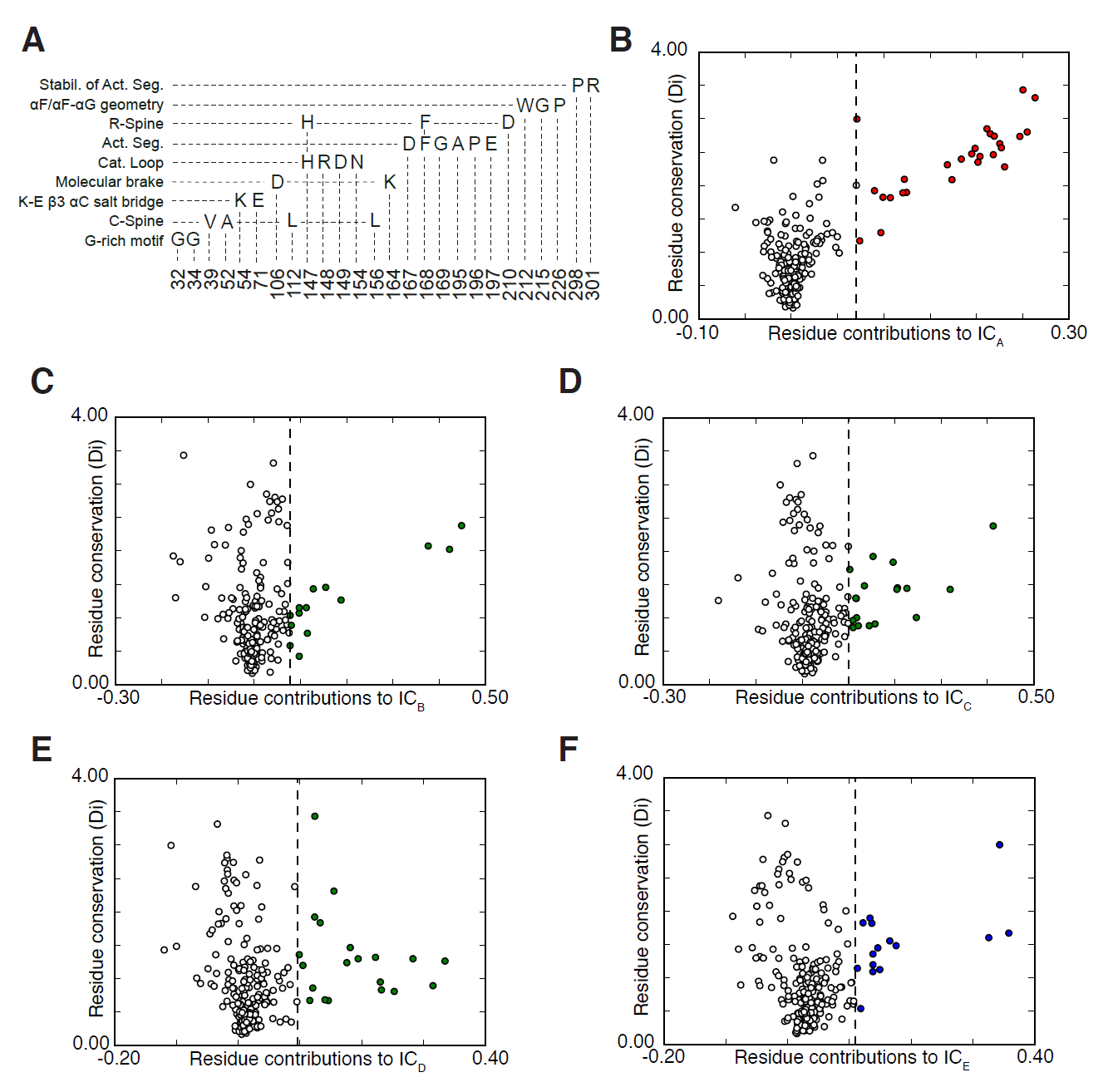
Description of the red sector and correlation between residue conservation and sector membership. A- Enumeration of the residues forming the red sector including their residue number in ERK2 (using numbering from PDB structure 4QTE) in the X axis and structural components that they are part of in the Y axis.
B- Scatterplots between residue conservation on the Y axis and residue contribution to the first independent component IC_A_. reproduced from Fig 2B, for comparison with panels BF. Residues forming the red sector are colored red.
C- Scatterplots between residue conservation on the Y axis and residue contribution to the second independent component IC_B_. Residues forming the green sector are colored green.
D- Scatterplots between residue conservation on the Y axis and residue contribution to the third independent component IC_C_. Residues forming the green sector are colored green.
E- Scatterplots between residue conservation on the Y axis and residue contribution to the fourth independent component IC_D_. Residues forming the green sector are colored green.
F- Scatterplots between residue conservation on the Y axis and residue contribution to the fifth independent component IC_E_. Residues forming the blue sector are colored blue.

**Figure S3.**
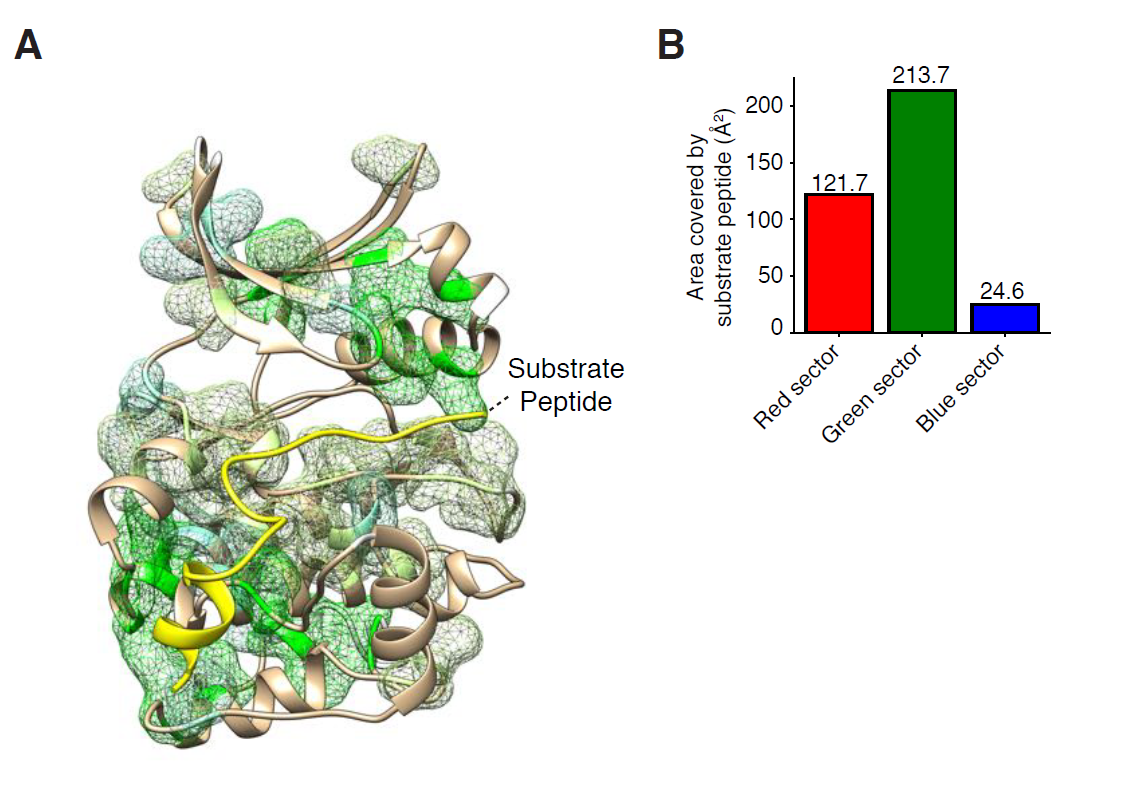
The green sector in the context of PKA in complex with PKI. A- Illustration of residues forming the green sector residues, superimposed on the structure of PKA in complex with PKI as a peptide inhibitor (PDB: 1ATP)^31^.
B- Quantification of the surface burial of red, green and blue sector residues by the peptide ligand., Solvent exposure in the presence and absence of the peptide was calculated using UCSF Chimera^32^.

**Figure S4.**
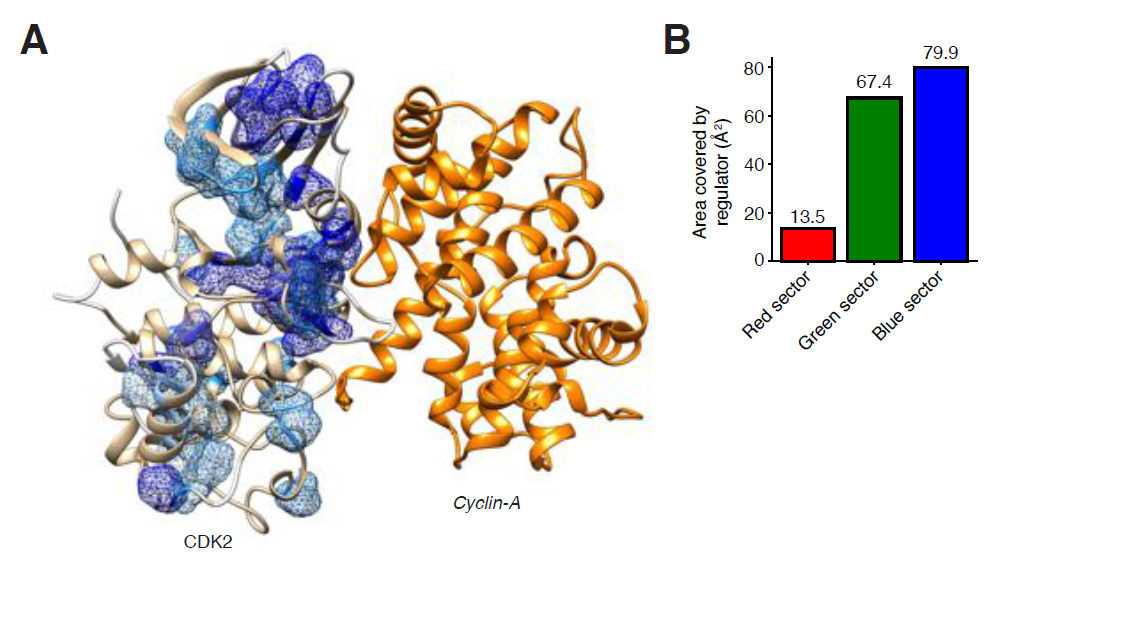
Blue sector residues in the context of CDK2 and its interaction with cyclin A. A- Illustration of residues forming the blue sector residues, superimposed on the structure of CDK2 in complex with cyclin-A (PDB: 1ATP)^31^.
B- Quantification of the surface burial of red, green and blue sector residues by the regulatory subunit, by calculating solvent exposure with UCSF Chimera^32^ in the presence and absence of cyclin-A (PDB: 1FIN).

**Figure S5.**
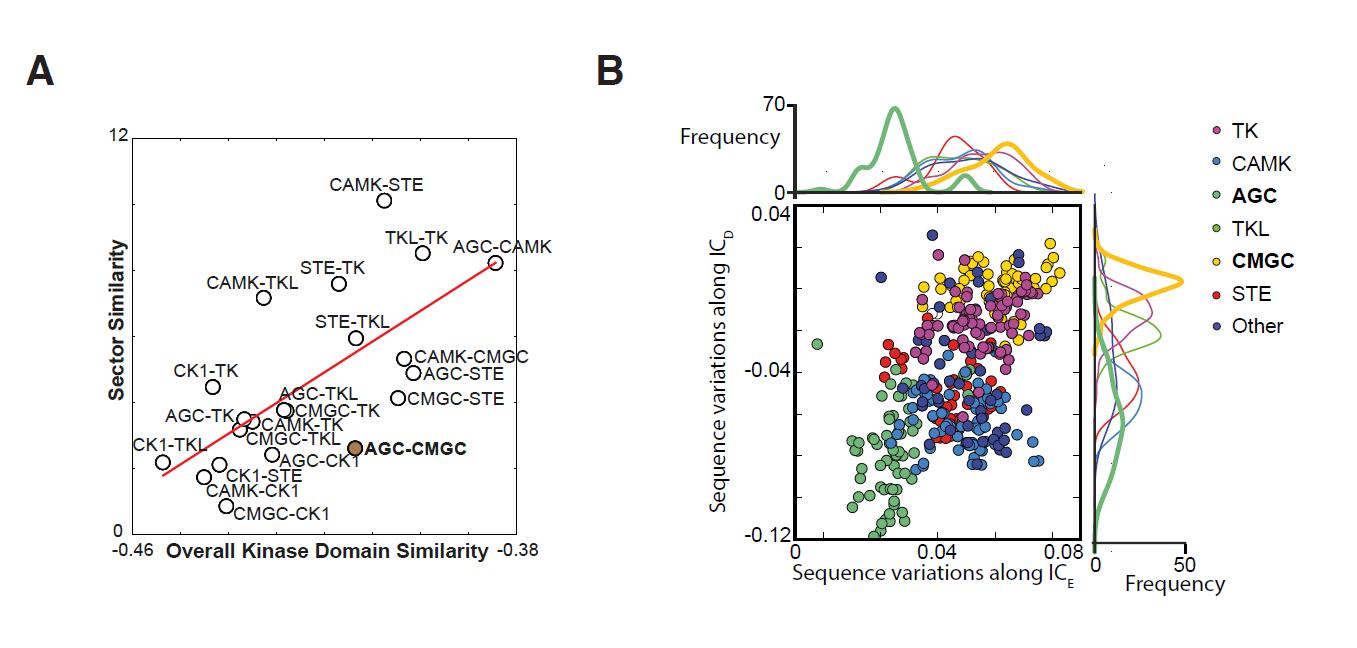
Sequence divergence of AGC and CMGC. A- Scatterplot from Figure 5A, annotated to indicate all pairs of kinase groups comparing their overall kinase domain similarity, as measured by overall BLOSUM distance^28^, in the X axis to degree of overlap in their sectors in the Y axis (see Methods). For completeness all kinase group pairs are labelled.
B- Scatterplot of sequence variations of residues that form the fourth and fifth independent components (IC_D_ and IC_E_) for all human protein kinases, compared to each other (see Methods). Kinases are colored by their kinase group. Note the large divergence between ACG (green) and CMGC (yellow) kinases.

### 5 Supplementary tables

**Table S1.** Number of kinases for the fifteen species and the seven well-established kinase groups used for our CCA models.

**Table S2.** Residues belonging to the different sectors in each of the seven representative kinases (and structures) used to describe each kinase group.

**Table S3.** Unique somatic cancer mutations mapped to the different human protein kinases with information regarding the sector membership of the site and corresponding mutation.

**Table S4.** Plasmids used in this study.

**Table S5.** Yeast strains used in this study.

## METHODS

### Statistical Coupling Analysis (SCA)

To perform SCA, we constructed an alignment including the kinase domain for protein kinases of all groups (kinome-wide alignment). Briefly, following alignment processing as detailed in previous work^11,12^, we used the python-based software package (pySCA) to compute a four-dimensional array with conservation-weighted covariance between all possible pairs of positions in the alignment and every possible amino acid residue within this pair of positions. By taking the magnitude (Frobenius norm) of the vector for all the amino-acids for a given pair of positions, this four-dimensional array was subsequently compressed into a two-dimensional coupling matrix, where the value at every position represents the degree of coupling between every pair of positions in the domain. Significantly coupled residues are identified by spectral decomposition and comparison to a randomized alignment, where residues within a kinase position are reshuffled thereby maintaining conservation while losing coupling. Next, independent component analysis (ICA) allows the identification of “independent components” (ICs) or clusters of residues that are more coupled amongst themselves than with the other residues. Positions contributing to each IC is defined by fitting an empirical statistical to the ICs and selecting positions above a defined default cutoff (>95% of the CDF). For further analysis of these independent components, by using singular value decomposition (SVD) as described in the next section, we can evaluate which specific protein sequences and domain positions contribute the most to a specific independent component^12^. Finally, as discussed elsewhere^12^, in cases where significant coupling still exist between several “independent components”, they are considered as part of the same protein sector.

### Singular Value Decomposition (SVD) and mapping of sequence variations along independent components

While a more complete theoretical description of Singular Value Decomposition (SVD) in the context of SCA can be found elsewhere^12^, here we provide a shorter description. Briefly, SVD allows to link coevolving groups of amino acid residues (such as those forming an independent component, IC, or protein sector) to patterns of sequence divergence in the original alignment. As such, using SVD we can map each protein in the original alignment as a function of its sequence divergence to every other protein in the alignment. Even more, by restricting the mapping to specific ICs the obtained mapping reflects the sequence relationship of each protein to every other protein specificifically as it relates to the amino acid residues forming that IC.

### Comparative Coupling Analysis (CCA)

Taking advantage of the seven standard kinase groups as classified on the basis of function, sequence and structural similarity, evolutionary history and broad substrate specificity (AGC, CAMK, CMGC, STE, CK1, TKL and TK)^2^, we constructed group-specific alignments by restricting each alignment to protein kinases belonging to that group. In order to allow the comparison between groups and with the kinome-wide alignment, the alignments were constructed with Mafft (with its parameters --add and --keeplength)^33^ using as baseline an original alignment including all human eukaryotic protein kinases. A canonical representative structure for each group was chosen based on completeness of the structure solved and optimizing for the largest number of residues within the kinase domain being covered. The canonical representative structures used were PKCtheta for AGC (PDB: 2JED), Pim1 for CAMK (PDB: 4JX3)^34^, TTBK1 for CK1 (PDB: 4BTJ)^35^, ERK2 for CMGC (PDB: 4QTE)^36^, MST2 for STE (PDB: 4LGD)^37^, BRaf for TKL (PDB: 4MBJ)^38^ and Abl for TK (PDB: 1FPU)^39^. Using each canonical representative structure and following the steps described in the SCA section above, we calculated coupling matrices for each kinase group separately. Once coupling matrices were calculated for the different kinase groups, they were mapped back to the representative structure that covered the largest number of residues within the kinase domain, namely ERK2 (PDB: 4QTE)^36^. By cross-comparing with the sectors identified in the kinome-wide analysis and other groups, we predicted in silico functional similarities and differences between the kinase groups. Finally, we quantified the degree to which a residue predicted to be of one sector in one kinase group tended to encode the same sector in other kinase groups and used the median number of groups encoding the same sector as a general measure of compositional conservation.

### Estimation of negative selection (Omega estimates)

Using the YN00 program that is part of the PAMLX package^40^ with default parameters we estimated the number of synonymous and nonsynonymous substitutions for the different residues to be considered, while correcting for multiple substitutions, transition/transversion rate biases and base/codon frequency biases^13^. These omega estimates are a measure of the amount of negative or positive selection that a specific protein or protein segment has gone through, with distributions around 1.0 indicating similar degrees of positive and negative selection and distributions below 1.0 indicating stronger negative selection). Four our purposes, after obtaining cDNA for all CMGC kinases from KinBase (kinase.com/kinbase), we constructed a cDNA alignment from the CMGC-specific alignment, allowing us to map back the sector sites that each codon corresponds to for a large number of sites, and computed omega estimates for the three different sectors as well as for non-sector sites.

### Residue conservation

The conservation of amino acid residues independently of other positions is here measured by the Kullback-Leibler relative entropy, Di. This measure compares the observed amino acid residue at a position to the background frequency of this amino acid from a non-redundant database of protein sequences.

### Calculation of area covered by substrate peptide or kinase regulator in kinase structures

The solvent exposure of every residue in the kinase domain is calculated using the UCSF Chimera package^32^ in the presence and absence of the substrate peptide or kinase regulator. In-house python scripts subsequently compare the solvent exposure calculated for both situations and calculate the area that is buried by the peptide or regulator in residues that form the red, green or blue sectors.

### Measuring similarity between kinase groups

Pairwise kinase group similarity was measured by calculating residue-normalized BLOSUM distances for every residue within the kinase domain as described elsewhere^17^. The coupling difference between two kinase groups is calculated by measuring the LogWorth, -log10(p-value), where the p-value is calculated from a hypergeometric test comparing the number of shared sector sites given the size of the sectors in both groups. As a result from these calculations, higher LogWorth values correspond to higher coupling similarity between kinase groups. After identifying AGC-CMGC as the pair of kinase most divergent in their coupling given their kinase similarity, their divergence is further inspected using SVD and other standard methods previously described^11,12^.

### Mapping of somatic cancer mutations

Genomic coordinates (human genome version GRCh38.p7) for missense cancer somatic point-mutations were retrieved from COSMIC v79^29^, and they were mapped to ENSEMBL canonical proteins, predicting the variants functional effect with the standalone perl script of the Ensembl Variant Effect Predictor, v87.18^41^. A total of 1,515,599 of cancer somatic mutations were mapped to a canonical protein. To obtain, for all protein kinases, the kinase domain residues perturbed by somatic cancer mutations, all the variants that mapped to the kinase domain were aggreagated by kinase residues. Only ENSEMBL canonical proteins with a 100% identical kinase domain sequence, with respect to the corresponding kinase domain sequence reported in KinBase, were considered further.

In order to define sectors for all protein kinases of all groups, using the kinome-wide alignment, the sector sites identified in the group representative kinases were mapped to the corresponding residues of all the other kinases within the groups. The kinase domain residues that in the kinome-wide alignment did not map to any residues of the corresponding group representatives, i.e. sequence insertions, due to the uncertainty in sector association, were excluded from the analysis. A total of 14,860 mutations were mapped to 13,152 sites within a kinase domain.

The mutation percentage was calculated across all kinases, for all sectors, as the number of residues perturbed by somatic cancer mutations, divided by the number of residues in the sector. Wilcoxon signed-rank tests were performed to assess the significance of the difference, across all kinases, between the mutation percentage in the red sector compared to the blue, the green and the non-sector. Definitions for OG and TSG were obtained from a work reviewing the functional role of the kinome in cancer^4^.

### Yeast strains and plasmids

Yeast strains and plasmids used in this work are described in Supplementary Tables 2 and 3, respectively. All strains are in the W303 genetic background. Gene deletions were performed by one-step PCR as described^42^. All mutants were integrated into yeast genome as a single copy expressed from their endogenous promoter.

### Site-directed mutagenesis

Site-directed mutagenesis was performed with QuickChange according to the manufacturer’s directions (Agilent).

### Cell growth and treatment with α factor

All cells were grown in synthetic complete media with dextrose (SDC). Three single colonies from each strain bearing the AGA1pr-YFP reporter were inoculated in 1 ml SDC in 2 ml 96-well deep well plates and serially diluted 1:5 three times. Plates were incubated overnight at 30°C. In the morning cells from the row that had been diluted 1:25 were typically found to have OD_600_ ˜0.5. These cells were diluted 1:5 in 4 rows of a 96 well U-bottom micro-titer plate in a total volume of 180 µl and incubated for 1 hour at 30°C. In each row, cells were treated with different concentrations of α factor: 0, 0.01, 0.1 and 1 µM (10x stocks of α factor were prepared and 20 µl were added to 180 µl cells). Treated cells were incubated for 4 hours at 30°C before translation was stopped by addition of 50 µg/ml cycloheximide. Cells were incubated for an additional hour at 30°C to allow time for fluorophores to mature.

### Flow cytometry

The AGA1pr-YFP reporter was measured by flow cytometry by sampling 10 µl of each sample using a BD LSRFortessa equipped with a 96-well plate high-throughput sampler. Data were left ungated and FlowJo was used to calculate median YFP fluorescence. Bar graphs show the average of the median of the three independent colonies that were assayed, and error bars are the standard deviation.

### Confocal microscopy

96 well glass bottom plates were coated with 100 µg/ml concanavalin A in water for 1 hour, washed three times with water and dried at room temperature. 80 µl of cells that had been treated with pheromone at the indicated concentrations for 3 hours were diluted to OD_600_ ˜0.05 and added to a coated well. Cells were allowed to settle and attach for 15 minutes, and unattached cells were removed and replaced with 80 µl SDC media. Imaging was performed at the W.M Keck Microscopy Facility at the Whitehead Institute using a Nikon Ti microscope equipped with a 100×, 1.49 NA objective lens, an Andor Revolution spinning disc confocal setup and an Andor EMCCD camera. Images were analyzed in ImageJ.

### Western blotting

Total protein was TCA purified from cells as described. 10 µl of each sample was loaded into 4-15% gradient SDS-PAGE gels (Bio-Rad). Gels were run at 25 mA for 45 minutes, and blotted to PVDF membrane at 225 mA for 40 minutes. After 1hr blocking in Li-Cor blocking buffer, membranes were incubated with anti-FLAG primary antibody (SIGMA, F3165) and/or anti-PGK (22C5D8) overnight at 4°C on a platform rotator (all 1:1000 dilutions in blocking buffer). Membranes were washed three times with TBST and probed by anti-mouse or anti-rabbit IR dye-conjugated IgG (Li-Cor, 926-32352, 1:10000 dilution). The fluorescent signal was detected with the Li-Cor/Odyssey system.

